# Respiratory Syncytial Virus Infections Induce Hypermetabolism in Pediatric Upper Airways

**DOI:** 10.1101/2020.07.18.200964

**Authors:** Svetlana Rezinciuc, Lavanya Bezavada, Azadeh Bahadoran, Jesse F. Ingels, Young-Yin Kim, Stephania A. Cormier, John P. Devincenzo, Barry L. Shulkin, Heather S. Smallwood

**Affiliations:** Department of Pediatrics, University of Tennessee Health Science Center, Memphis, Tennessee, USA; Department of Genetics, Genomics, and Informatics, University of Tennessee Health Science Center, Memphis, Tennessee, USA; Department of Biological Sciences, Louisiana State University, and Department of Comparative Biomedical Sciences, Louisiana State University School of Veterinary Medicine, Baton Rouge Louisiana, USA; Department of Diagnostic Imaging, St. Jude Children’s Research Hospital, Memphis, Tennessee, USA

**Keywords:** RSV (respiratory syncytial virus), Metabolism, pediatric, Glycolysis, OXPHOS = oxidative phosphorylation, viral infection, Bioenergetics, Metabolomics, Infant, virus, infection, respiratory infection

## Abstract

To determine whether respiratory syncytial virus (RSV) regulates human metabolism, we used positron emission tomography (PET) of patient lungs along with bioenergetics and metabolomics of patient upper airway cells and fluids. We previously found a significant negative monotonic relationship between glucose uptake and respiratory viral infection in 20 pediatric patients (e.g., 70% of infected patients had glucose uptake within 0–3 days). In our recent study, 3 out of 4 patients positive for glucose uptake at later times (>5 days) were positive for RSV infection. At present, the bioenergetics of upper respiratory cells (URCs) from nasal pharyngeal aspirates have not been investigated, and *in vitro* studies indicate RSV reduces metabolism in cell lines. To define metabolic changes in RSV-infected pediatric patients, we acquired fresh aspirates from 6 pediatric patients. Immediately following aspiration of URCs, we measured the two major energy pathways using an XFe flux analyzer. Glycolysis and mitochondrial respiration were significantly increased in URCs from RSV-infected patients, and mitochondrial respiration was operating at near maximal levels, resulting in loss of cellular capacity to increase respiration with impaired coupling efficiency. Metabolomics analysis of metabolites flushed from the upper airways confirmed a significant increase in TCA cycle intermediates. Taken together, these studies demonstrate RSV induces significant hypermetabolism in pediatric patients’ lungs and respiratory tract. Thus, hypermetabolism is a potential anti-viral drug target and reveals RSV can regulate human metabolism.

**Contributions to the field:** Metabolic changes in humans in response to viral infection are largely unknown. In this brief clinical report, we find metabolism is markedly increased in live upper respiratory cells from infants infected with respiratory syncytial virus (RSV) concomitant to changes in metabolites in their upper airway fluids. This sheds light on viral induced hypermetabolism in the airways and offers potential biomarkers for RSV. In addition, this identifies potential therapeutic targets for host directed therapies of aberrant metabolism in RSV. This work has clinical impact as biomarkers and therapeutics for RSV are needed for this pervasive virus that causes infections with long term consequence for some children. Further, advancements in molecular mechanisms underpinning RSV infection biology are constrained by the difficulties in translating model systems to humans as well as relating human studies in adults to infants (Mestas and Hughes, 2004; Papin et al., 2013).

## 1 Introduction

Respiratory syncytial virus (RSV) infects 70% of infants by 12 months and reemerges as a serious lower respiratory tract illness in the elderly. RSV treatment is mostly supportive care because vaccines or therapies are unavailable (Oshansky et al., 2009; Hurwitz, 2011; Simoes et al., 2015; Heylen et al., 2017). Although RSV’s immediate effects can be catastrophic, up to 50% of hospitalized children develop long-term complications persisting into adulthood (Hall et al., 2013). RSV etiology and exacerbation have been attributed to host genetic factors (Miyairi and DeVincenzo, 2008; Grad et al., 2014), innate and adaptive immune responses (Kurt-Jones et al., 2000; Haynes et al., 2001; Murawski et al., 2009; You et al., 2013; Cormier et al., 2014; Huang et al., 2015; Schmidt and Varga, 2017), pathophysiological factors (Becnel et al., 2005), and an immature immune system with delayed adaptive immune responses (Derscheid and Ackermann, 2013). The pulmonary innate immune response is the first-line defense. The molecular mechanisms of epithelial and immune responses in RSV-infected children are needed to implement prevention, identify biomarkers, and find therapeutics. However, advancements in RSV infection biology are limited due to difficulties in translating *in vitro* and *in vivo* models to humans as well as relating human studies in adults to infants (Mestas and Hughes, 2004; Papin et al., 2013).

Every cell produces reactive oxygen species (ROS) in mitochondria. ROS participate in cell signaling and increase with elevated metabolism. RSV and influenza increase ROS production in epithelial cells and some immune cells upon infection. Indeed, oxidative stress markers were elevated in nasopharyngeal secretions and blood from RSV-infected children (Hosakote et al., 2011). Specific inhibitors of complexes in mitochondrial respiration blocked RANTES (chemotactic factor) in RSV-infected A549 cells, indicating rapid mitochondrial ROS generation (<2 hr post infection) (Garofalo et al., 2013). RSV is also implicated in dysregulation of cellular metabolic homeostasis (Oshansky et al., 2009; Cervantes-Ortiz et al., 2016). Additionally, RSV significantly reduced bioenergetics in A549 and MH-S cell lines (Grunwell et al., 2018; Hu et al., 2019). Although these studies were in cell lines, they demonstrate involvement of mitochondrial metabolism in RSV infection. Nevertheless, cell lines inaccurately reflect respiratory-induced changes in host metabolism. Primary human respiratory cells are required to target host metabolism during respiratory antiviral drug identification (Smallwood et al., 2017).

Here, we quantified glycolysis and mitochondrial respiration of epithelial and immune cells isolated from nasopharyngeal aspirates (NPAs) from naturally infected, non-ventilated pediatric patients. We validated metabolic changes with metabolomics of upper respiratory fluids and identified six metabolites significantly altered by RSV. Since our retrospective study of positron emission tomography (PET) scans indicated RSV-induced hypermetabolism in pediatric patients, we determined if RSV induced hypermetabolism in the respiratory tract and altered metabolic pathways. This information will advance our understanding of RSV-host interactions for developing metabolite-targeting drugs (Smallwood et al., 2017). Nasal lavage fluids can be collected noninvasively from infants and children and provide temporary relief by clearing the sinuses. Thus, we determined if metabolites are sufficiently abundant in nasal lavage fluids for detection via mass spectrometry (MS) and identification of RSV biomarkers.

## 2 Materials and Methods

### Subjects and study procedures

Inclusion criteria required participants who met the clinical case definition of RSV infection or were asymptomatic. This study was conducted in compliance with 45 CFR46 and the Declaration of Helsinki. Institutional Review Boards of the University of Tennessee Health Science Center/Le Bonheur Children’s Hospital approved the study. Participants provided nasal swabs and nasal lavages after enrollment. Parents ranked participants’ symptom severity and duration. St. Jude Children’s Research Hospital approved the retrospective study of respiratory infected patients who received PET scans.

### Infected patient PET scans

These previously published data and methods were reanalyzed with respect to RSV. Briefly, patients with normal glucose levels received I.V. injections of fluorodeoxyglucose (FDG) after fasting. Relaxed, prone patients remained in a quiet, dark room. One hour later, transmission computed tomography (CT) and PET images were captured with a GE Discovery LS PET/CT system or 690 PET/CT system (GE Medical Systems, Waukesha, WI). Vendor-supplied software was used for reconstruction, and standardized uptake values were determined.

### Pediatric nasal pharyngeal aspirates

After enrollment subjects were swabbed and nasal rinses obtained. Nasal aspirates were obtained at enrollment, placed on ice immediately and small aliquot removed for diagnostics. Clinical diagnostics were performed including antigen test and quantitative reverse transcription PCR (RT-qPCR). The aspirates were immediately transported on ice to the research laboratory and cell separated by gently centrifuging. The supernatant was stored at -80°C, and the viability and number of upper respiratory cells (URCs) were determined.

### Upper respiratory cell bioenergetics

URCs (200,000 per well) were immediately seeded in XFe96 plates following the manufacturer’s cell suspension protocol. The glycolytic stress test and mitochondrial stress test were performed in separate wells in the same plate to expedite measurements. Four to eight wells of technical replicates were run per patient sample. DNA per well was quantified with CyQUANT (Thermo Scientific, Waltham, MA) and used for data normalization. Data analysis was performed using Agilent Seahorse Wave software v2.6.1 (Agilent, Santa Clara, CA).

### Single Cell RNA sequencing (scRNA-Seq)

URCs were thawed and washed in HBSS with BSA, counted and enzymatically treated to reduce the mucosity. 5-10,000 cells per subject were filtered to remove dead cells, fixed with DSP, then 800 cells captured and single cell mRNA prepared for sequencing using Fluidigm C1 coupled to Ti2 Inverted imaging system with NIS Elements software (Nikon). scRNA-seq libraries of full length polyA-positive mRNA’s were generated for each cell using SMART-Seq v4 technology (Takara). For barcoding, each C1-HT plate was divided into 20 columns of 40 cells each and each well labeled with a position specific barcode and each column was given a separate Nextera XT i7 index (Illumina). The resulting 800 cDNA’s were pooled and NEBNext multiplex oligos for Illumina (the i5 indexes; New England BioLabs) was used as a dual index primer. Ten C1 plates were combined for analysis using the NovaSeq 6000 System (Illumina). We used SingleR to cluster by cell types per subject and we analyzed enrichment of KEGG pathways per cell type.

### Metabolomics of upper respiratory fluids

We performed metabolite extraction and UPLC–HR mass spectral analysis. Briefly, metabolites were solvent extracted, solvent evaporated, resuspended in water, and placed in a chilled autosampler for mass spectrometric analysis. Aliquots (10 µL) were injected through a Synergi 2.5 micron reverse-phase Hydro-RP 100, 100 x 2.00 mm LC column (Phenomenex, Torrance, CA) and introduced into the MS via an electrospray ionization source conjoined to an Exactive™ Plus Orbitrap Mass Spectrometer (Thermo Scientific). We used full-scan mode with negative ionization mode (85–1000 m/z), 3 kV spray voltage, 10 psi flow rate at 320°C, 3e6acquisition gain control, 140,000 resolution with scan windows of 0 to 9 minutes at 85 to 800 m/z and 9 to 25 minutes at 110 to 1000 m/z and solvent gradient (Lu et al., 2010). Data files generated by Xcalibur (Thermo Scientific) were converted to open-source mzML format using ProteoWizard (Martens et al., 2011, Chambers et al. 2012). Maven (mzRoll) software (Apache Software Foundation, Wakefield, MA) automatically corrected total ion chromatograms based on the retention times for each sample and selected unknown peaks (Clasquin et al. 2012; Melamud et al. 2010). Metabolites were manually identified and integrated using known masses (± 5 ppm mass tolerance) and retention times (Δ ≤ 1.5 min).

Multivariate statistical analysis for MS/MS data was performed using XLSTAT OMICS (Addinsoft, New York, NY) with Excel (Microsoft Corporation, Redmond, WA). To ensure observations were directly comparable and to account for respiratory secretion concentrations, peak intensity was normalized to total intensities. These data were independently k-means clustered followed by ascendant hierarchical clustering based on Euclidian distances. Data values of the permuted matrix were replaced by corresponding color intensities based on interquartile range with a color scale of red to green through black. Unsupervised multivariate principal component analysis (PCA) was performed, and the difference in metabolite concentrations per group was determined using one-way ANOVA with Benjamini-Hochberg post hoc correction. Significant differences were detected using Tukey’s honest significant difference (HSD) test for multiple comparisons. Mean intensity data and standard deviation for each metabolite were graphed in Prism (GraphPad, San Diego, CA) and tested for significance using unpaired t-test.

## 3 Results

### Hypermetabolism in the lungs of pediatric patients diagnosed with respiratory infections

We performed a retrospective study of pediatric patients diagnosed with respiratory viral infections confirmed by RT-qPCR who underwent FDG-PET/CT (Smallwood et al., 2017). FDG uptake is proportional to the metabolic rate of a region, and hypermetabolic lesions, regions, and foci are readily detected with FDG-PET/CT (Kostakoglu et al., 2003; Jadvar et al., 2005; Sharp et al., 2008; 2011; Davis et al., 2018). Several patients showed hypermetabolism in tumor-free lungs (Smallwood et al., 2017). We also found a significant temporal relationship: the sooner patients were scanned after infection diagnosis, the more likely they were to have hypermetabolic regions in their lungs (Smallwood et al., 2017). Representative images of uninfected, RSV infected, and a subject we followed for

6 month as the RSV induced hypermetabolic regions subsided [Fig 1 A, B, and C respectively]. In these studies, we grouped all respiratory viruses together including RSV, metapneumovirus, RSV and metapneumovirus co-infection, adenovirus, parainfluenza, or influenza. Recently, we compared the temporal distribution of subjects separated by virus. Patients with glucose uptake were unevenly distributed among respiratory pathogens. Compared with influenza-infected patients (black + bars), RSV-infected patients were positive for glucose uptake in the lungs (red + bars) much longer **[Fig 1D]**. Additionally, 3 out of 4 patients scanned one week after RSV diagnosis exhibited FDG uptake in their lungs.

**Figure 1.**
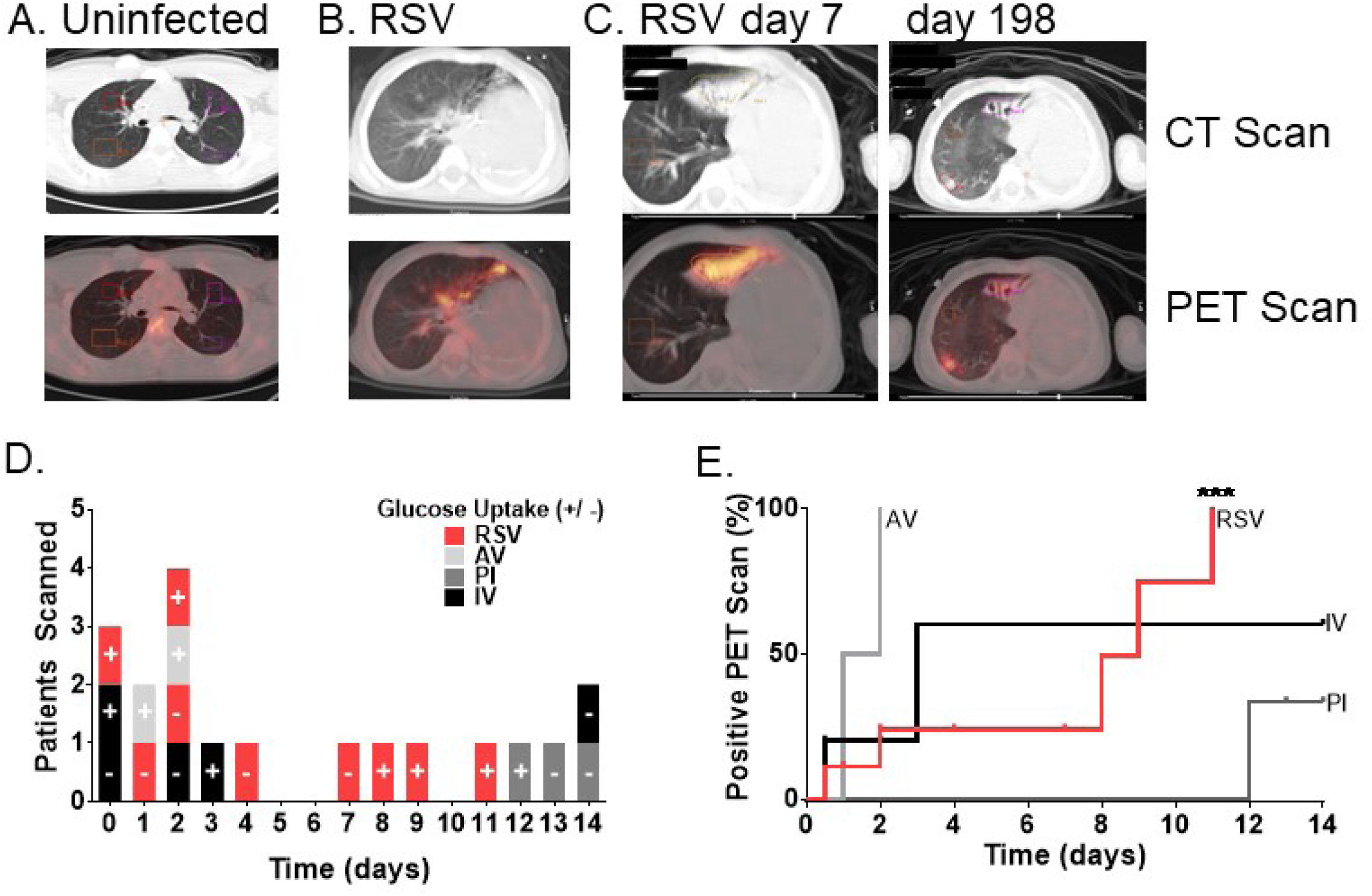
Hypermetabolism in the lungs of pediatric patients diagnosed with respiratory infections. We performed a retrospective study of pediatric patients who received PET scans within 15 days of clinical diagnosis with respiratory viral infection. Acceptance criteria included confirmation of clinical diagnosis with RT-qPCR within 15 days of PET scan. Whole body transmission CT and PET images were obtained after patients with normal blood glucose fasted for 4 or more hours and were given 5.5 MBq/kg FDG intravenously followed by a 1-hr uptake period. (A) The number of patients scanned is plotted against time in days from RT-qPCR to PET scan, and zero indicates the scan and RT-qPCR were performed on the same day. Subjects are colored by viral group with positive or negative symbols indicating presence or absence of glucose uptake in the PET scan. (B) The Kaplan and Meier product limit method was used to create curves for the infected subjects at risk for hypermetabolism diagnosed by PET scan, and the curves were compared with log-rank tests (AV: Adenovirus; PI: Parainfluenza virus; IV: Influenza virus). The Mantel-Cox log-rank test and the Gehan-Breslow-Wilcoxon test indicated the curves were significantly different with p-values of 0.0018 and 0.0038, respectively. The Pearson correlation test was performed on each event risk curve. Both PI and RSV infected groups had significant temporal correlations (associated p-values of 0.0078 and 0.0250, respectively) represented by asterisks. With Pearson’s r values of -0.9922 and -0.8165 and R2 = 0.9845 and 0.6667, respectively.

We performed time-to-event analysis on these groups to determine the proportion of patients who likely had hypermetabolic regions due to respiratory viral infections. We plotted the number of subjects at risk over time from RT-qPCR to PET scan using the product limit estimator method (Kaplan and Meier) to estimate the proportion of infected individuals who likely had glucose uptake in their lungs within 2 weeks of diagnosis. Adenovirus-infected patients had a median of 1.5 days, and influenza-infected patients had 3 days. By contrast, RSV had a median event time of 9 days. These curves were significantly different by log-rank using the Mantel-Cox test **[Fig 1E]**. We used the Pearson correlation test to determine the relationship between glucose uptake risk (percent) and time elapsed between diagnosis by RT-qPCR and PET scan. RSV had a significant positive relationship (r value: 0.9117, R^2^: 0.8312, p value: 0.0006), indicating hypermetabolism in RSV-infected patients’ lungs continued throughout the study interval, with higher associated risk one week after diagnosis.

To determine if these metabolic changes occurred in normal children with community-acquired RSV infections, we obtained NPAs from 11 hospitalized pediatric patients. Pediatric patients, neither intubated nor admitted to the ICU, were enrolled and swabbed for RSV antigen testing followed by NPA collection (Oshansky et al., 2014). We determined cell viability and numbers with acridine orange and propidium iodide. Five patients were excluded due to low cell viability and numbers. Six patients, all under two years old, were included (Table 1). Following the initial screening for RSV antigen, RSV A and RSV B levels were quantified by RT-qPCR (Table 1). One RSV antigen-negative patient was RT-qPCR positive and moved to the infected group. A small level near baseline was detected in one control. However, based on the lack of RSV symptoms and testing negative for RSV antigen, this subject remained in the control group.

**Table 1.** Pediatric participant anthropomorphic data, nasal swab results (antigen test), NPA cell viability, NPA cell number, and RT-qPCR results.

| Age (days) | Symptom Duration (days) | Antigen Test | Viability (%) | Cell Number (per ml) | RSV A RT-qPCR (PFUe/ml) | RSV B RT-qPCR (PFUe/ml) | Subject Identifier |
| --- | --- | --- | --- | --- | --- | --- | --- |
| 142 | 3 | - | 71 | 1.45 x 10<sup>7</sup> | 0 | 0.15 | Ctl 1<sup>‡</sup> |
| 415* | 4 | - | 67 | 1.59 x 10<sup>6</sup> | 0 | 0 | Ctl 2 |
| 457* | 7 | - | 74 | 5.61 x 10<sup>6</sup> | 0 | 5.93 | RSV 1<sup>‡</sup> |
| 57 | 3 | + | 84 | 2.13 x 10<sup>7</sup> | 0 | 5 | RSV 2<sup>‡</sup> |
| 149 | 21 | + | 77 | 1.57 x 10<sup>7</sup> | 6.39 | 0 | RSV 3 |
| 117 | 7 | + | 77 | 3.44 x 10<sup>6</sup> | 0 | 4.13 | RSV 4<sup>‡</sup> |

\* Adjusted age corrected for premature birth; <sup>‡</sup> Negative for influenza

### RSV increases glycolysis in upper airway cells

URCs were immediately plated for bioenergetic analysis on an Xfe96 bioanalyzer. Bioenergetic states are based on substrate consumption for ATP production, and product efflux varies with cellular metabolism. Glycolysis tightly correlates with extracellular lactic acid accumulation (i.e. R^2^ = 0.9101), and lactic acid excretion per unit time in glycolysis accounts for most pH changes in most cell types (Legmann et al., 2011; TeSlaa and Teitell, 2014). Thus, Xfe96 measures the extracellular acidification rate (ECAR) as a proxy for glycolysis. However, infected immune cells use radical generation and pH in signaling and innate effector functions. Therefore, we used the glycolytic stress test to distinguish ECAR from glycolytic lactate from other cellular acidification sources (TeSlaa and Teitell, 2014; Zhou et al., 2015; Thomas, 2017). ECAR was quantified after glucose was added to URCs, followed by inhibiting ATP synthase with oligomycin to maximize glycolysis and hexokinase with 2-deoxyglucose (2-DG) **[Fig. 2A]**. 2-DG completely blocks glycolysis, allowing quantification of ECAR independent of glycolysis. After establishing baseline, we added glucose and determined basal ECAR. To isolate glycolytic ECAR, we subtracted residual non-glycolytic ECAR from basal ECAR. RSV infection significantly increased glycolysis in pediatric URCs **[Fig 2A]**. We then determined the maximal glycolytic output of URCs by inhibiting oxidative phosphorylation (OXPHOS) of ADP to ATP and the electron transport chain with oligomycin to force URCs to use glycolysis for ATP production. URCs from RSV-infected patients doubled their glycolytic capacity **[Fig 2A]**. The glycolytic reserve is the difference between maximal glycolytic output and basal glycolysis and reflects the potential of cells to further increase reliance on glycolysis to meet energy demands. Glycolytic reserve significantly increased with RSV infection [**Fig 2A]**. The final injection of 2-DG allowed quantification of non-glycolytic acidification, which significantly increased in RSV-infected patients, nearly two-fold greater than that in uninfected controls **[Fig 2A]**.

**Figure 2.**
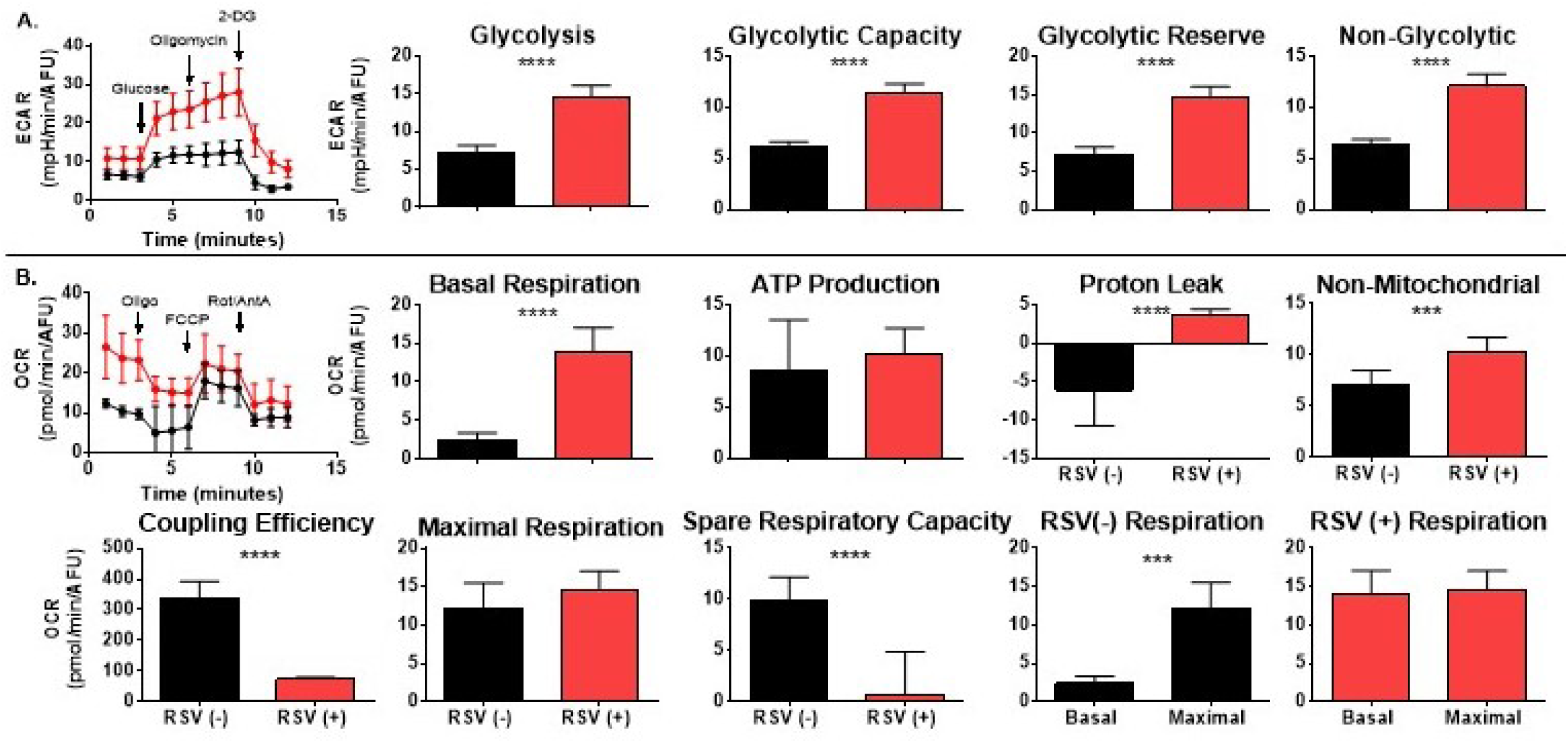
RSV induced significant increases in glycolysis and mitochondrial respiration. After NPA collection and enumeration, 200,000 viable URCs per well were plated in technical replicates distributed on one plate per patient. Each plate was subjected to the Glycolysis Stress Test and Cell Mito Stress Test in parallel on the same plate. The plates and cells were processed and read on XFe96 following the manufacturer’s protocols. Bioenergetics data processing and analysis of XFe assays was done using XF96 software. Oxygen consumption rates (OCRs) and respiratory parameters and extracellular acidification (ECAR) and glycolytic parameters were derived from kinetic rate data calculated using stress test profiles per manufacturer’s guidelines. These data were exported to Prism. Bar graphs represent the mean and standard deviation of each patient group with asterisks symbolizing p-values determined using two-tailed unpaired t tests (P values: 0.05 (*), <0.01 (**), <0.001 (***) and ≤ 0.0001 (****)).

### RSV increases mitochondrial respiration in upper airway cells

To determine the flux of pyruvate into the TCA cycle to fuel OXPHOS, we used respirometry with Xfe96. We quantified the oxygen consumption rate (OCR) of URCs and isolated mitochondrial respiration with the mitochondrial stress test **[Fig 2B]**. The underlying principle of this test is the amount of oxygen consumed during respiration is stoichiometrically related to the amount of ADP and substrate of respiratory oxidation. Sequential inhibition of complex V (ATP synthase) and complexes I and III of the electron transport chain with oligomycin and a combination of rotenone and antimycin A, respectively, allow determination of respiration efficiency. OCR from non-mitochondrial sources such as NADPH oxidases can be quantified following complete inhibition of the mitochondrial electron transport chain by the final inhibitor, rotenone/antimycin A. We can then determine and deduct the non-mitochondrial OCR, which can be high in activated immune cells.

ATP fluctuations control basal mitochondrial respiration, which relies on and oscillates with substrate availability. Minor changes in maximal respiratory capacity or proton leak have little impact on basal respiration. Nonetheless, cell size or number can affect basal respiration. Thus, we used equal numbers of URCs and calculated the respiration rate per amount of DNA. RSV infection dramatically increased basal respiration approximate 5.5 times that of uninfected controls **[Fig 2B]**. After defining basal respiration, we isolated the rate of mitochondrial ATP synthesis by adding oligomycin and quantifying the corresponding decrease in respiration. ATP production remained unchanged with increased proton leak following RSV infection **[Fig 2B]**. However, oligomycin slightly hyperpolarizes mitochondria, so this assay may underestimate ATP synthesis by less than 10% (Brand and Nicholls, 2011). Although RSV-infected URCs had significantly higher glycolysis, they maintained the ability to increase glycolysis in response to oligomycin, requisite for quantifying ATP production with this assay. After oligomycin, non-mitochondrial OCRs can be subtracted, and correction for hyperpolarization by oligomycin can help determine proton leak. RSV infection increased proton leak in URCs **[Fig 2B]**. Proton leak values for uninfected URCs are a function of inherently high non-mitochondrial oxygen consumption in this mixed cell population, including a large population of monocytes (Huang et al., 2015).

In OXPHOS, substrate oxidation is coupled to ADP phosphorylation to ATP while mitochondria establish proton-motive force by pumping out protons that are returned by ATP synthase, thereby generating ATP. Mitochondrial coupling efficiency varies with ATP demand, is sensitive to mitochondrial dysfunction, and is similar to the phosphate/oxygen ratio (Brand and Nicholls, 2011). Mitochondrial coupling efficiency is ATP production divided by proton leak. RSV-infected URCs had significantly reduced mitochondrial coupling efficiency **[Fig2B]**. Next, we uncoupled oxygen consumption from ATP production by adding the protonophore FCCP to determine the maximal respiratory capacity of URCs. We found no difference between infected and uninfected maximal respiration **[Fig 2B]**. Notably, this method indirectly measures maximal mitochondrial respiration, representing maximal mitochondrial respiration within cellular substrate uptake and metabolism that influence respiratory chain activity.

Calculating spare respiratory capacity by subtracting basal respiration from the maximal respiration rate can determine how closely cells operate near their OXPHOS threshold. Spare respiratory capacity is a cellular bioenergetic diagnostic because it reflects the cell’s ability to coordinate substrate supply and oxidation with electron transport in response to increased energy demand (Nicholls et al., 2007; Yadava and Nicholls, 2007; Choi et al., 2009). RSV-infected URCs had significantly decreased spare respiratory capacity, amounting to a nearly 8-fold reduction **[Fig 2B]**.

These bioenergetics analysis were on bulk URCs. To determine the contribution of cell subsets, we performed single cell transcriptomics on these revived URCs. We used SingleR to cluster by cell types per subject and we analyzed enrichment of KEGG pathways per cell type. We found CD8+ T cells, monocytes and neutrophils were enriched in several pathways associated with viral replication and infection (e.g. RNA transport and HIV/HCV/EBV infection) are upregulated following RSV infection **[Fig 3A-C]**. We found epithelial cells in fresh URCs, with viral inclusion bodies, using microscopy (DNS). Yet we did not find epithelial cells with scRNA-seq and suspected these cells were depleted by our removal of dead cells prior to cell capture. Indeed, when we performed flow cytometry on banked URCs samples from RSV infected children, we found immune cells but very few live epithelial cells (DNS). A similar study used the quantitative proteomic analysis identified significant enrichment of immune cell pathways in revived RSV-infected URCs (Aljabr et al., 2019). Thus, we anticipated the enrichment of these pathways but were surprised to find relatively low metabolic pathway changes in revived URC immune cells. It is possible epithelial cells are the main contributors to the significant increase in metabolism in RSV infected URCs, but we cannot rule out the effects of cryopreservation on dampening immunometabolism. Thus, it appears similar to bioenergetics, single cell transcriptomics may require fresh URCs to capture the metabolic response of URCs.

**Figure 3.**
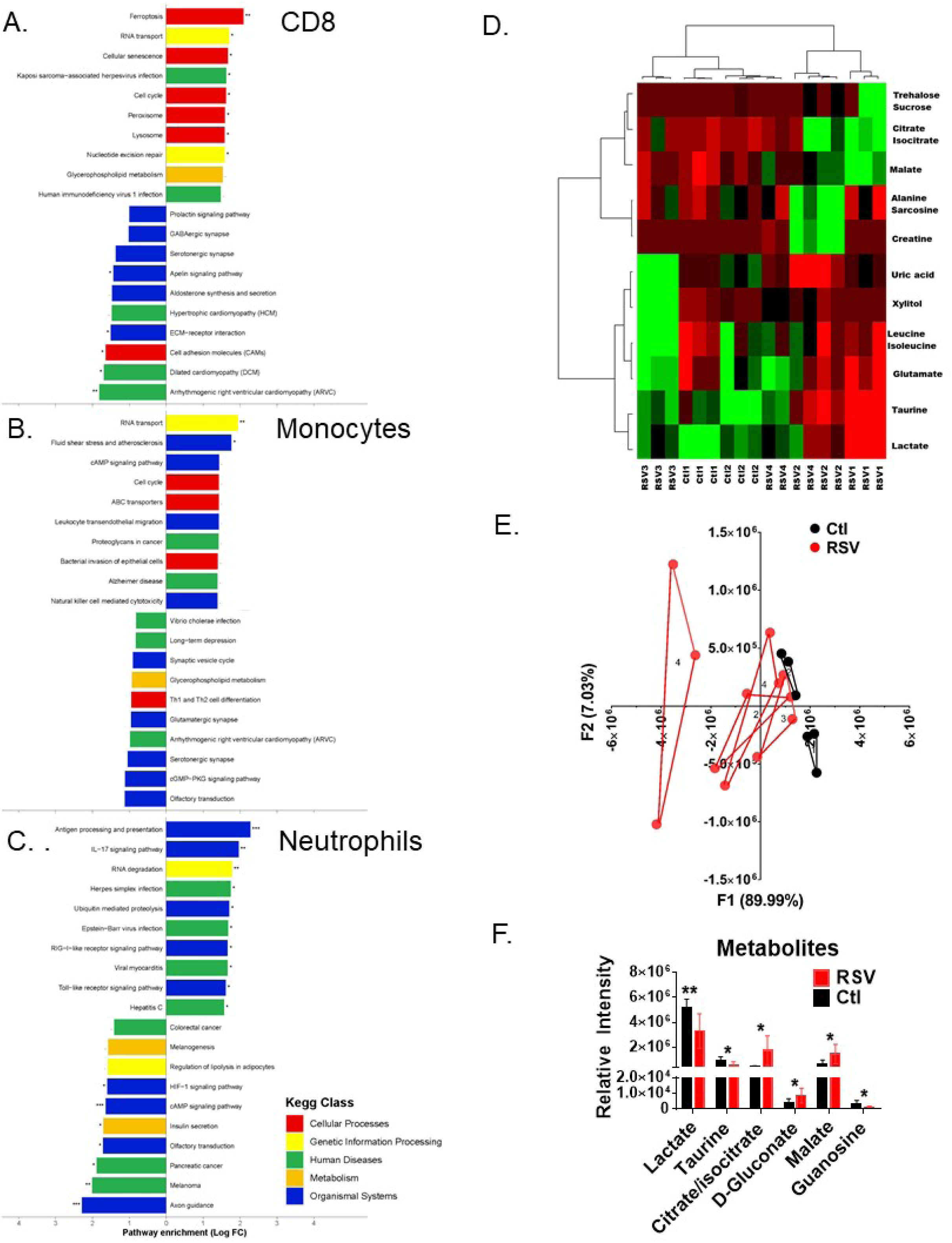
RSV infection changes metabolite composition in upper respiratory fluids. URCs were thawed, washed, and filtered to remove dead cells. 5-10,000 cells per subject were fixed, 800 cells captured and single cell mRNA prepared for sequencing using Fluidigm C1 platform. scRNA-Seq libraries of full length polyA-positive mRNA’s were generated for each cell using SMART-Seq v4 technology (Takara). For barcoding, each C1-HT plate was divided into 20 columns of 40 cells each and each well labeled with a position specific barcode and each column was given a separate Nextera XT i7 index (Illumina). The resulting 800 cDNA’s were pooled and given a dual index primer (NEBNext multiplex oligos for Illumina). Ten C1 plates were combined for analysis using the NovaSeq 6000 System (Illumina). We used SingleR to cluster by cell types per subject and we analyzed enrichment of KEGG pathways per cell type (A-C). Metabolites were extracted from 50µl of NPA supernatant and subjected to UPLC–HRMS metabolomics. Metabolites were extracted from 50 µl NPA supernatant and subjected to UPLC–HRMS metabolomics analysis three times per sample. Metabolites were manually identified and integrated using known masses (± 5 ppm mass tolerance) and retention times (Δ ≤ 1.5 min). Peak intensity was normalized to the sum of peak intensity. (A) Metabolites were K-means clustered followed by ascendant hierarchical clustering based on Euclidian distances and nondynamic metabolites excluded (0.25 < std dev). Metabolite clusters were also represented via dendrograms displayed vertically for metabolites and horizontally for patients. The data values of the permuted matrix were replaced by corresponding color intensities based on interquartile range with color scale of red to green through black, resulting in a heat map. (C) Unsupervised multivariate principal component analysis (PCA) was performed, resulting in F1 and F2 with a cumulative percent variability of 97.02%. Replicated for each subject are joined by lines and subject number indicated in the triangle center. (E) One-way ANOVA with Tukey’s honestly significant difference test and Benjamini-Hochberg post hoc correction was used to identify metabolites with significant differences among the patient groups (XLSTAT, Addinsoft). Relative intensity data for each metabolite were graphed in Prism. Bar graphs represent the mean and standard deviation of each patient group with asterisks symbolizing p-values determined using two-tailed unpaired t tests [P values: 0.05 (*) and <0.01 (**)] (F).

### RSV infection alters metabolites in upper airway fluids

Next, we wanted to determine if airway fluids reflect the molecular mechanisms driving metabolic changes in RSV-infected URCs. To identify any differences in metabolites in upper airway fluids from RSV-infected patients, we performed targeted discovery metabolomics (TDM) using MS. After cells were removed for bioenergetic measures, we organically extracted metabolites from the NPA supernatant. Metabolites were injected through a reverse-phase LC column and analyzed on an Exactive™ Plus Orbitrap Mass Spectrometer (Thermo Scientific). The spectra were taken in full-scan mode with negative ionization (Lu et al., 2010). RAW files were converted to open-source mzML format, and total ion chromatograms for each sample were corrected based on retention times (Melamud et al., 2010; Martens et al., 2011; Chambers et al., 2012). We previously determined the masses (± 5 ppm mass tolerance) and retention times (Δ ≤ 1.5 min) of 300 metabolites associated with disease pathologies. We manually identified and integrated NPA metabolites using our previously determined masses and retention times for targeted discovery. With TDM-MS, we identified 35 metabolites in the NPA [Sup File 1].

To determine the dataset structure and relationships between groups, we used unsupervised multivariate statistical analysis. Metabolites and individuals were clustered independently using k-means clustering followed by ascendant hierarchical clustering based on Euclidian distances. We arranged data matrices according to the clustering with spatial relationships proportional to similarity among patients or metabolites **[Fig 3D]**. The top horizontal dendrogram separates patients while the vertical groups metabolites, each by how strongly their concentrations correlate. With metabolite peak intensities rearranged according to metabolite clustering, the metabolites are divided into two distinct groups on the y-axis **[Fig 3D-left dendrogram]**. These groups show metabolite concentrations depending on RSV infection. The patient samples are also divided into two groups **[Fig 3D-top dendrogram]**. Interestingly, RSV-infected patient 3 (RSV3) grouped with the healthy control (Ctl), whereas RSV patients 1, 2, and 4 clustered together **[Fig 3D]**. When we decoded the samples, patient 3 had been symptomatic for 21 days, while the other RSV-infected patients were symptomatic for a week or less **[Table 1]**. This finding might reflect the kinetics of returning to metabolic homeostasis in the upper airway. However, more data are needed to confirm this idea. The control and RSV3 group display an inverse pattern of metabolites with a relatively low concentration in the top metabolite cluster and relatively high concentrations in the bottom cluster **[Fig 3D]**. In contrast, RSV patients 1, 2, and 4 exhibit lower concentrations for most metabolites.

To evaluate group trends and sample uniformity as well as identify potential outliers, we used PCA. Variation was explained by F1 and F2 very well with a cumulative percent variability of 95.57% **[Fig 3E]**. Furthermore, the F1 or F2 squared cosines of each patient observation exceeded 0.5 (DNS). Again, RSV patient 3 had a closer relationship to controls than to patients with more recent symptom onset **[Fig 3E (red triangle 3)]**. The overlap of RSV patient 3 and controls likely reflects the trajectory of this patient’s recovery **[Table 1]**. Accordingly, patient 3 was removed from comparative analysis for RSV-associated metabolites.

To identify specific metabolic products that change during RSV infection, we compared uninfected controls to RSV-infected patients who were symptomatic for less than a week (acute RSV). We used one-way ANOVA with Tukey’s HSD test and Benjamini-Hochberg post hoc correction to identify metabolites with significant differences among the groups. Six upper respiratory fluid metabolites were significantly different in patients with acute RSV **[Fig 3F]**. Significant depleted compounds in RSV infected samples included lactate, taurine, and guanosine. On the other hand, RSV infection triggered increased levels of some metabolites including citrate, D-gluconate and malate compared to non-infected samples.

## Discussion

Our retrospective study of pediatric patients who received PET scans within 15 days of clinical diagnosis with respiratory viral infection showed RSV-infected patients are positive for glucose uptake in the lungs much longer than influenza-infected patients [Fig 1A]. To determine if these metabolic changes were reflected in children with community-acquired RSV infections, we performed bioenergetic analysis on URCs obtained from pediatric patients diagnosed with RSV. We found RSV infection significantly increases glycolysis in pediatric URCs. Furthermore, glycolytic reserve significantly increased with RSV infection **[Fig 2A]**. By contrast, *in vitro* influenza infection had no effect on normal human bronchial epithelial cells nor glycolytic reserve in murine dendritic cells (Smallwood et al., 2017; Rezinciuc et al., 2020). Altogether, these observations indicate URCs in RSV-infected pediatric patients RSV are highly glycolytic. Accumulating evidence suggests viruses induce glycolysis to facilitate viral replication (Sanchez and Lagunoff, 2015; Yu et al., 2018; Passalacqua et al., 2019).

OCR quantification in fresh URCs indicates RSV significantly increased basal respiration and proton leak without affecting ATP production. Significantly reduced mitochondrial coupling efficiency was also observed **[Fig 2B]**. When comparing cells, coupling efficiency is a reliable indicator of mitochondrial dysfunction as an internally normalized ratio sensitive to changes in all bioenergetic flux components (Brand and Nicholls, 2011). With increased basal respiration and unaffected ATP production, proton re-entry is the major respiratory influence, and severely blunted maximal mitochondrial respiration suggests RSV-induced changes in respiration occur at substrate oxidation upstream of the proton circuit (Hafner and Brand, 1991; Porter and Brand, 1995) (Brand and Nicholls, 2011) Nonetheless, declining maximum respiratory capacity is considered a strong indicator of potential mitochondrial dysfunction. URCs from RSV-infected children show a significant drop in spare respiratory capacity, demonstrating how closely cells are operating at their bioenergetic threshold. In RSV infection, induced basal respiration is nearly maximized. Loss of spare respiratory capacity also indicates mitochondrial dysfunction, and this loss of function “may not be particularly apparent under basal conditions, when respiration rate is strongly controlled by ATP turnover, but becomes manifested only under load when ATP demand increases and substrate oxidation more strongly limits respiration” (Brand and Nicholls, 2011) Indeed, this issue became apparent when comparing the basal and maximal respiration of uninfected patients’ URCs, revealing a significant difference **[Fig 2B, RSV(-) Respiration]**. In contrast, basal and maximal respiration rates were nearly identical in RSV-infected patients’ URCs **[Fig 2B, RSV (+) Respiration]**. These data indicate RSV infection increases respiration in URCs to near maximal levels, resulting in loss of spare respiratory capacity with impaired coupling efficiency and increased proton leak, ultimately detrimental to efficient ATP production.

MS analysis of metabolic products changing during RSV infection shows increased lactate in upper airway fluid. Consistently, glycolysis significantly increased in RSV-infected patient cells. However, lactate turnover flux exceeds that of glucose and is the highest of all circulating metabolites (Annison et al., 1963; Okajima et al., 1981; Hui et al., 2017) Consistent with RSV dramatically increasing mitochondrial metabolism, ^13^C-lactate extensively labels TCA cycle intermediates equal to glucose in all tissues except the brain and is the primary contributor to tissue TCA metabolism in fasting. In addition to increased URC mitochondrial metabolism with significantly reduced airway fluid lactate levels, TCA cycle intermediates citrate and malate were significantly increased in the upper respiratory fluids of RSV-infected patients [Fig 3C]. In the lung, citrate and malate concentrations are higher than other TCA intermediates like succinate and are TCA products derived from glucose and lactate metabolism in the lung (Hui et al., 2017).

In most cells, glucose is metabolized via glycolytic pyruvate in the TCA cycle, which efficiently produces ATP (Vander Heiden et al., 2009). In contrast, most cancers and transformed cell lines use aerobic glycolysis, referred to as the Warburg effect, where the demand for anabolic metabolites drives less efficient ATP production by uncoupling glycolysis and the TCA cycle to allow carbon commitment to macromolecule production as opposed to ATP output regardless of the availability of oxygen. This phenotype reflects an increase in metabolic demands for cell proliferation (Vander Heiden et al., 2009). Our metabolic and bioenergetic studies show that RSV re-wires metabolism in a manner that resembles Warburg metabolism. However, we found a very significant increase in basal respiration in the URC and a significant decrease in lactate in the upper respiratory fluids. Typically, one would expect a decrease in OCR and increase in lactate accumulation for traditional Warburg metabolism, but more studies are required to precisely define carbon metabolism in RSV infection. Further, if RSV infection aims to rapidly change metabolism in a proviral phenotype, kinetics may be more important that ATP efficiency. Indeed, in some studies glycolysis supports flexible carbon metabolism with rapid production of ATP (i.e. faster than the complete oxidation of glucose in the mitochondria). However, we measured the ATP production supported through respiration not glycolysis. We did not recover enough URCs to perform an ATP rate assay in parallel, but our future studies will dissect the contribution of both pathways. As noted above, the mitochondrial stress these underestimates ATP production. How this reconciles with high glycolysis and mitochondrial respiration as measured here remains to be seen, but it is apparent RSV infected cell bioenergetics are very high irrespective of efficiency of ATP production. We found it surprising that both glycolysis and mitochondrial respiration were significantly increased, even if respiration efficiency is compromised by RSV infection, ATP is still being produced at least as well as uninfected. Glycolysis, in the Warburg metabolic phenotype discussed above, actually produces enough ATP for genome and cell proliferation in cancer. Here on wonders if the RSV infected cells unusually high metabolism contributes to rapid cell death.

Our data indicate community-acquired RSV infection dramatically increases glycolysis and mitochondrial metabolism in the airways of pediatric patients. We found strong evidence of hyperglycolytic metabolism in patient lungs and significant increased glycolysis and mitochondrial respiration in respiratory cells and fluids from the upper respiratory tract of infected patients. However, our patient studies contradict current findings delineating decreased bioenergetics using RSV infection of A549 cells and MH-S cell lines (Grunwell et al., 2018; Hu et al., 2019) and a more recent study by Martín-Vicente et al 2020 that indicated Warburg (i.e. increase glycolysis with decreased TCA) in A549 following RSV. Why these *in vitro* models contradict our patient findings remains unknown, but based on our previous studies with influenza it is likely due to the inherent metabolic nature of transformed cell lines (Smallwood et al., 2017). Clearly, URCs show significantly increased glycolysis and mitochondrial respiration reflected by the metabolites flushed out with upper airway fluids. These findings may be instrumental in developing an accurate cell model to study RSV-induced changes in host metabolism of the respiratory tract.

The bioenergetics of patient URCs remain undefined in RSV or other respiratory infections. The absence of progress is most likely due to the bioenergetic instability of repository cells following rescue from cryopreservation. Indeed, metabolic recovery of URCs from freeze thaw overwhelms effects induced by infection (DNS). To avoid the effect of freeze thawing, we collected NPAs from six patients and immediately measured their bioenergetics. In the future, we will perform single cell transcriptomics on fresh URCs to capture epithelial cells. These studies are limited by patient number because measuring bioenergetics in delicate URCs from our repository is challenging. Nonetheless, with fresh URCs, we confirmed previous results: RSV increased bioenergetics in the lungs of adult and neonatal C57BL/6 mice based on oxygen consumption and ATP concentrations (Alsuwaidi et al., 2014). After unsuccessfully sorting and measuring URC bioenergetics, these analyses were performed on bulk URCs. Without single cell metabolomics, we cannot define the relative contribution of cell subsets within URCs to changes in bioenergetics. We are currently addressing this issue and are aware these findings represent initial clinical observations. Although incomplete, the evidence obtained from our samples suggests community-acquired RSV has a profound effect on respiratory metabolism, representing a potential drug target.

## Acknowledgments

We thank the staff of Le Bonheur Children’s and St Jude Children’s Research Hospitals for their work caring for their patients and supporting our studies. We thank Lisa Harrison and Elizabeth Meals for enrolling and consenting subjects and sample collection and the Le Bonheur Children’s Hospital molecular diagnostics lab. Mass spectrometric analysis was performed at the Biological and Small Molecule Mass Spectrometry Core, University of Tennessee, Knoxville, TN, with the assistance of Dr. Shawn R. Campagna, Dr. Hector F. Castro, Sara Howard and Eric Tague.

## Author Contributions

S.R. and L.B NPA metabolite and cell sample preparation, drug/cell titrations, XFe96 assays; A.B. data analysis and writing; S.R., J.F.I. sc-RNA-Seq; Y.Y.K. RT-qPCR; S.C. URC characterization; J.P.D. clinical coordination, enrollment, and sample collection; B.L.S. retrospective clinical study and PET scan analysis; H.S.S. data analysis, experiment coordination, and manuscript writing.

## Ethics statements

### Studies involving animal subjects

#### Generated Statement

No animal studies are presented in this manuscript.

### Studies involving human subjects

#### Generated Statement

The studies involving human participants were reviewed and approved by UTHSC Institutional Review Board (IRB). Written informed consent to participate in this study was provided by the participants’ legal guardian/next of kin.

### Inclusion of identifiable human data

#### Generated Statement

No potentially identifiable human images or data is presented in this study.

### Data availability statement

#### Generated Statement

The datasets generated for this study are available on request to the corresponding author.

